# *Escherichia coli* NusG links the lead ribosome with the transcription elongation complex

**DOI:** 10.1101/871962

**Authors:** Robert S. Washburn, Philipp K. Zuber, Ming Sun, Yaser Hashem, Bingxin Shen, Wen Li, Sho Harvey, Stefan H. Knauer, Joachim Frank, Max E. Gottesman

## Abstract

It has been known for more than 50 years that transcription and translation are physically coupled in bacteria, but whether or not this coupling may be mediated by the two-domain protein N-utilization substance (Nus) G in *Escherichia coli* is still heavily debated. Here, we combine integrative structural biology and functional analyses to provide conclusive evidence that NusG can physically link transcription with translation by contacting both RNA polymerase and the ribosome. We present a cryo-electron microscopy structure of a NusG:70S ribosome complex and nuclear magnetic resonance spectroscopy data revealing simultaneous binding of NusG to RNAP and the intact 70S ribosome, providing the first direct structural evidence for NusG-mediated coupling. Furthermore, *in vivo* reporter assays show that recruitment of NusG occurs late in transcription and strongly depends on translation. Thus, our data suggest that coupling occurs initially via direct RNAP:ribosome contacts and is then mediated by NusG.

## Introduction

Gene expression is a universal process in all cells and consists of transcription, i.e. the synthesis of RNA based on the DNA, and – if RNA is not the final gene product – translation, i.e. the messenger RNA (mRNA)-guided synthesis of a protein. Since the late 1960s it has been known that the rates of transcription and translation are synchronized in *Echerichia coli* so that mRNA is translated while being transcribed (Das et al., 1967; Mehdi and Yudkin, 1967; Miller et al., 1970; Proshkin et al., 2010; Vogel and Jensen, 1994, 1995). This process, called transcription:translation coupling, is possible due to the lack of a physical barrier between transcription and translation in bacteria (reviewed in (Conn et al., 2019)). Only recently, direct physical interactions between RNA polymerase (RNAP) and the ribosomes have been demonstrated (Demo et al., 2017; Fan et al., 2017; Kohler et al., 2017), consistent with earlier observations that transcriptional events may control translation activity and *vice versa*(Proshkin et al., 2010). As transcription and translation are closely connected to other central processes in a bacterial cell, such as DNA repair (Pani and Nudler, 2017) and protein folding (Thommen et al., 2017), transcription:translation coupling constitutes one of the key resulatory functions in bacterial gene expression.

However, there are also indications that transcription:translation coupling may involve a member of the family of N-utilization substance (Nus) G proteins, which serves as adapter connecting RNAP and the lead ribosome (Burmann et al., 2010, 2012; Saxena et al., 2018; Zuber et al., 2019). *E. coli* Nus G, member and eponym of the only universally conserved class of transcription factors (Werner, 2012), consists of two domain, an N- and a C-terminal domain (NTD and CTD, resepectively) connected v*ia* a flexible linker, which move independently (Burmann et al., 2011; Mooney et al., 2009a). NusG-NTD binds RNAP and accelerates transcription elongation (Burova et al., 1995; Kang et al., 2018; Mooney et al., 2009a). Structural studies demonstrate that NusG-CTD, which is a five-stranded β-barrel with a Kyrpides-Ouzounis-Woese motif (Kyrpides et al., 1996), is a versatile binding platform for different transcription factors. By binding to protein S10, which is part of the 30S subunit of the ribosome, NusG may link transcription and translation (Burmann et al., 2010). Saxena *et al* also demonstrated specific 1:1 binding of NusG to 70S ribosomes both *in vitro* and *in vivo* (Saxena et al., 2018).

S10 is identical with transcription factor NusE and forms a ribosome-free complex with NusB, NusA and NusG which suppresses transcription termination (Dudenhoeffer et al., 2019; Krupp et al., 2019; Said et al., 2017; Squires et al., 1993). Finally, NusG-CTD binds to termination factor Rho and is required for most Rho activity *in vivo* (Burmann et al., 2010; Lawson et al., 2018; Mitra et al., 2017). Transcription:translation coupling prevents Rho factor from terminating transcription by sequestering the NusG-CTD and by blocking Rho access to RNAP via untranslated mRNA. Cryptic *E. coli* Rho-dependent terminators located within open reading frames (orfs) are revealed when ribosomes are released by polar nonsense mutations (Cardinale et al., 2008; Newton et al., 1965).

Nevertheless, there is evidence for intragenic uncoupling and Rho-dependent transcription termination in the absence of nonsense mutations. Washburn and Gottesman (Washburn and Gottesman, 2011) and Dutta et al. (Dutta et al., 2011) found that Rho resolves clashes between transcription and replication. Such conflicts are likely to occur within, rather than at the end of, genes. Uncoupling would allow Rho to release the stationary transcription elongation complexes (TECs).

Mutations in *nusE* or *nusG* that uncouple transcription from translation increase sensitivity to chloramphenicol (Saxena et al., 2018). This antibiotic retards translation, breaking the bond between the lead ribosome and TEC. Uncoupled TEC may backtrack or terminate prematurely (Dutta et al., 2011).

In this report, we present a cryo-electron microscopy (cryo-EM) structure showing NusG binding to the S10 subunit in a 70S ribosome. The NusG-CTD binding site of S10 is also target of the ribosome-release factor, transfer-messenger (tmRNA), raising the possibility that tmRNA might displace NusG at rare codons, thereby uncoupling transcription from translation (Roche and Sauer, 1999). We also show by solution-state nuclear magnetic resonance (NMR) spectroscopy that NusG, once bound to RNAP, can interact with S10 or with a complete ribosome, setting the structural basis for coupling.

NusG couples transcription with translation *in vivo*, as proposed earlier (Burmann et al., 2010). Uncoupling of RNAP from the lead ribosome is enhanced when translation is compromised. Importantly, we demonstrate that uncoupled RNAP can outpace translation, leading to Rho-dependent transcription termination. This intragenic termination explains the necessity for the apparent perfect synchronization between transcription and translation (Proshkin et al., 2010).

## Results

### Structural evidence of NusG binding to the ribosomal S10 subunit on a 70S ribosome

We assembled a NusG:70S complex by incubating 70S ribosomes with an excess of NusG and determined the structure of this complex by cryo-EM and single-particle reconstruction. Overall, 188,127 particles were extracted from 1327 images and ∼5% of these particles showed an extra mass of density attached to the mass identified as protein S10 (Fig. 1A,B). This additional density perfectly matches the size of NusG-CTD, suggesting that NusG binds at the site predicted from the solution NMR structure of NusG-CTD bound to the free ribosomal protein S10 in a 1:1 stoichiometry (Fig. 1A,B; (Burmann et al., 2010)). The density map reconstructed from the class of NusG:70S particles was refined to an average resolution of 6.8 Å. No density could be observed for NusG-NTD, indicating that it is flexibly bound to the NusG-CTD and does not interact with the ribosome.

**Figure 1.**
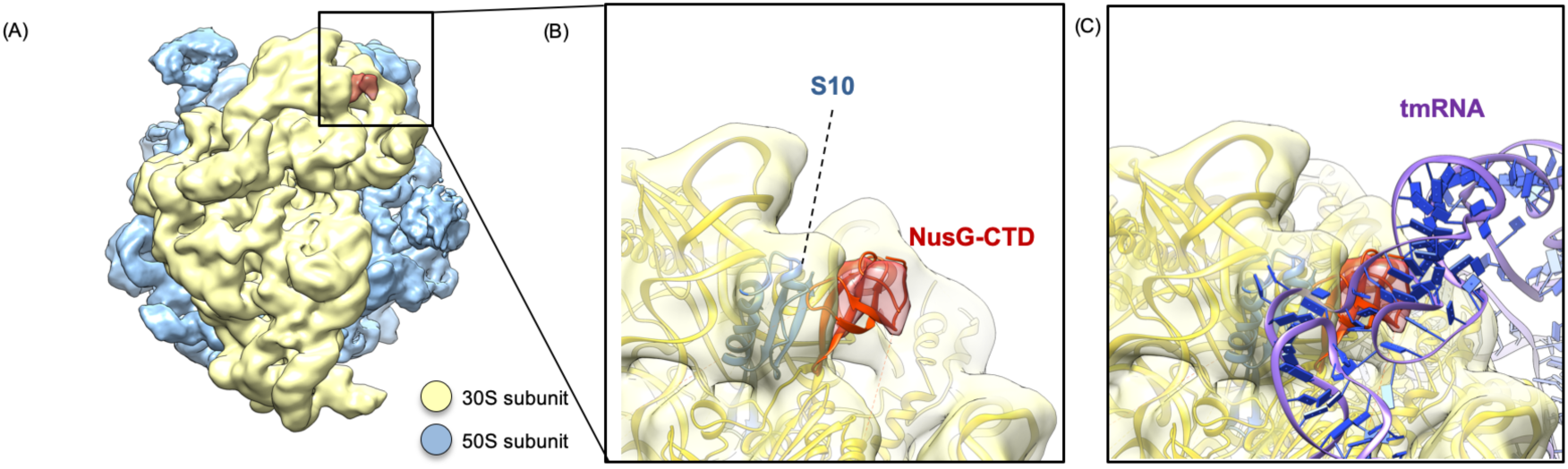
Structure of NusG-CTD bound to 70S ribosome. (A) Cryo-EM density of the 70S ribosome:NusG complex. The density of the 50S subunit is shown in light blue, the density of the 30S subunit in yellow, the density corresponding to NusG-CTD in red. **(B)** Close-up view of the region boxed in **(A)**. 70S (yellow), S10 (blue), and NusG-CTD (red) are in ribbon representation, cryo-EM density is shown as transparencies. **(C)** Superposition of the 70S:NusG complex with the 70S:tmRNA complex (tmRNA is in ribbon representation, purple and dark blue; EMD 5234, PDB: 3IZ4). 30S and NusG-CTD are displayed as in **(B)**.

During translation ribosomes may stall on incomplete mRNAs, i.e. they reach the 3’ end of an mRNA without terminating, resulting in an unproductive translation complex. Together with the small protein B (SmpB) tmRNA can bind to these stalled ribosomes in order to rescue them and to tag the nascent polypeptide chain for degradation in a process called trans-translation (Weis et al., 2010). Interestingly, the NusG-CTD binding site overlaps with the region of S10 that is contacted by the tmRNA when it is bound to a ribosome in its resume state (Fig. 1C; (Burmann et al., 2010; Fu et al., 2010; Rae et al., 2019; Weis et al., 2010)). From this we conclude that NusG-CTD and tmRNA share binding sites on S10, raising the possibility that, in addition to releasing stalled ribosomes, tmRNA competes with NusG for ribosome binding, thus preventing NusG from maintaining a linkage between the lead ribosome and RNAP. In other words, tmRNA might be able to displace NusG and thereby facilitate uncoupled transcription.

### Simultaneous binding of NusG to S10 and RNAP

In the cryo-EM structure of *E. coli* NusG bound to a paused TEC (Kang et al., 2018) only the density of NusG-NTD was observable, indicating that NusG-CTD moves freely and does not interact with RNAP. Binding of NusG-CTD to S10 was observed both in a binary system (Burmann et al., 2010) and a λN-dependent antitermination complex (Krupp et al., 2019; Said et al., 2017).

Since the NusG-CTD:S10 interaction is a prerequisite for NusG-mediated transcription:translation coupling, we probed this contact when NusG was bound to RNAP - but not in an antitermination context - by solution-state NMR spectroscopy. We employed NusG samples where [^1^H,^13^C]-labeled methyl groups of Ile, Leu, and Val residues in perdeuterated proteins served as NMR-active probes ([ILV]-NusG) to increase sensitivity, allowing us to study large systems.

In the methyl-transverse relaxation optimized spectroscopy (methyl-TROSY) spectrum of free [ILV]-NusG (Fig. 2A), signals of the NusG-NTD and NusG-CTD perfectly superimpose with the signals of the isolated [ILV]-labeled protein domains, suggesting that the domains move independently, confirming a previous report stating that there are no intramolecular domain interactions (Burmann et al., 2011). Upon addition of RNAP in a two-fold molar excess, [ILV]-NusG signals were significantly decreased in the one-dimensional methyl-TROSY spectrum (Fig. 2B, inset), indicating [ILV]-NusG:RNAP complex formation. Binding of RNAP increases the molecular mass of [ILV]-NusG dramatically, resulting in enhanced relaxation, which ultimately leads to drastic line broadening and a decrease in signal intensity. Interestingly, the two-dimensional spectra revealed a non-uniform signal decrease (Fig. 2B), which is caused by a combination of several effects. First, there is a general loss of signal intensity due to the increase in molecular mass upon complex formation, as discussed above. Second, upon binding, methyl groups of Ile, Leu, and Val residues located in the binding surface come into close proximity of RNAP protons. Dipole-dipole interactions contribute to relaxation processes so that the signal intensity of these methyl groups is decreased more strongly than that of methyl groups located elsewhere in [ILV]-NusG. Finally, signal intensities may be affected by chemical exchange processes. We analyzed the signal intensity of [ILV]-NusG signals in the presence of RNAP quantitatively by calculating relative signal intensities, i.e. the ratio of the remaining signal intensity of [ILV]-NusG in the presence of RNAP to the signal intensity of free [ILV]-NusG (Figure 2-figure supplement 1).

**Figure 2.**
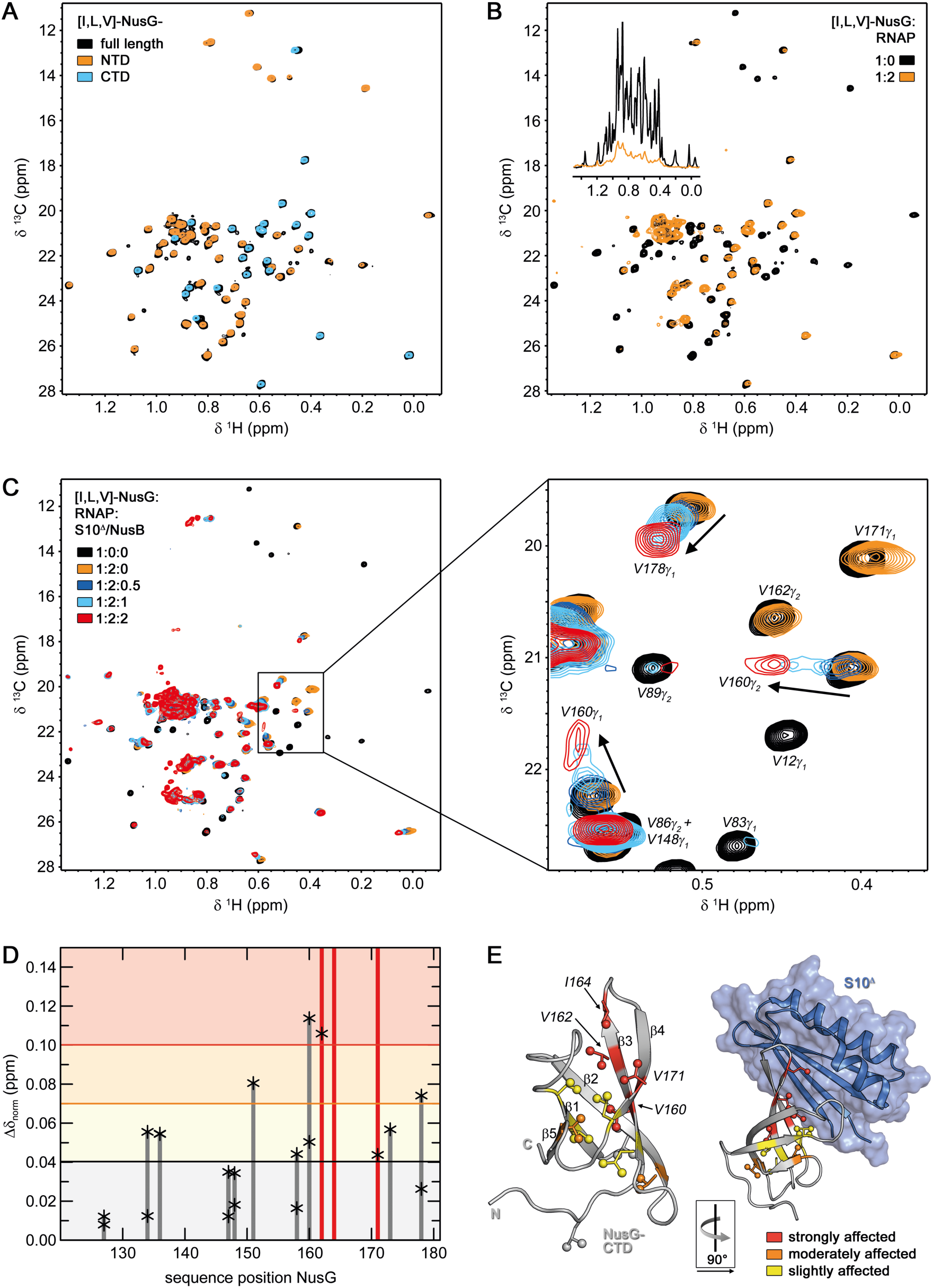
RNAP-bound NusG interacts with S10. (A) Superposition of 2D [^1^H, ^13^C]-methyl-TROSY spectra of [ILV]-NusG (black, 20 µM), [ILV]-NusG-NTD (orange, 100µM), and [ILV]-NusG-CTD (cyan, 30 µM). **(B)** 2D [^1^H, ^13^C]-methyl-TROSY spectra of [ILV]-NusG in the absence (black, 20 µM) and presence (orange, 18 µM) of two equivalents of RNAP. Inset: Normalized 1D [^1^H,^13^C]-methyl TROSY spectra, colored as 2D spectra. **(C)** 2D [^1^H, ^13^C]-methyl-TROSY spectra of [ILV]-NusG alone (20 µM), in the presence of a twofold molar excess of RNAP (18 µM [ILV]-NusG), and upon titration of [ILV]-NusG:RNAP with 218 µM S10^Δ^:NusB. The molar ratio of [ILV]-NusG:RNAP:S10^Δ^:NusB is indicated in color. The panel on the right shows an enlargement of the boxed region. Selected signals are labeled and arrows indicate chemical shift changes upon S10^Δ^:NusB addition. **(D) [**^1^H,^13^C]-methyl-TROSY-derived normalized chemical shift changes of [ILV]-NusG-CTD methyl group signals of RNAP-bound [ILV]-NusG upon complex formation with S10^Δ^:NusB. Asterisks mark the values of individual methyl group signals, bars represent the highest values. Red bars indicate vanishing signals. Horizontal lines are thresholds for affected methyl groups: slightly affected (0.04 ppm ≤ Δδ_norm_ < 0.07 ppm; black), moderately affected (0.07 ppm ≤ Δδ_norm_ < 0.1 ppm; orange), and strongly affected (Δδ_norm_ ≥ 0.10 ppm; red). **(E)** Mapping of affected methyl groups on the structure of isolated NusG-CTD (left; PDB ID: 2JVV) and NusG-CTD in complex with S10^Δ^(right; PDB ID 2KVQ). NusG-CTD is shown in ribbon (gray), S10^Δ^ in ribbon and surface (blue) representation. Affected Ile, Leu, and Val residues are colored according to **(D),** non-affected Ile, Leu, and Val residues are gray. Side chains of Ile, Leu, and Val residues are depicted as sticks, their methyl groups as spheres. Strongly affected Ile, Leu, and Val residues are labeled. The orientation of NusG-CTD in the complex relative to the isolated state is indicated.

The average relative intensity of NusG-NTD signals was significantly lower than that of the linker or the NusG-CTD, suggesting that NusG-NTD binds to RNAP whereas NusG-CTD remains flexible and moves independently, able to interact with other partners, as indicated by the NusG:TEC structure (Kang et al., 2018). The signal intensity of all Ile, Leu, and Val residues in the RNAP binding site of NusG was completely extinguished, confirming that NusG-NTD binds to RNAP at its known binding site (Drögemüller et al., 2015; Kang et al., 2018; Krupp et al., 2019; Said et al., 2017).

To test if NusG-CTD can bind to S10 while being tethered to RNAP *via* NusG-NTD, we titrated the [ILV]-NusG:RNAP complex with S10^Δ^ (Fig. 2C). In order to increase stability, we used this S10 variant lacking the ribosome binding loop in complex with NusB (Luo et al., 2008). Chemical shift changes of [ILV]-NusG-CTD signals upon titration of [ILV]-NusG:RNAP with S10^Δ^:NusB were determined (Fig. 2D) and affected residues were mapped onto the three-dimensional structure of NusG-CTD (Fig. 2E). Strongly affected residues are found to be located in β-strands 3 and 4 as well as in the connecting loop, in agreement with the binding site observed in the binary NusG-CTD:S10^Δ^ complex (Burmann et al., 2010). The loop between β-strands 1 and 2 is also part of the NusG-CTD:S10^Δ^ binding site, but as it does not contain any Ile, Leu, or Val residues, no NMR-active probes are available in this region; nevertheless, affected residues can be found in β-strand 1, directly preceding this loop. This suggests that the CTD:S10^Δ^ binding surface in the RNAP:NusG:S10^Δ^:NusB complex is identical to the one determined in the binary system. Importantly, the NusG-NTD signals do not change when S10 is added to the NusG:RNAP complex, indicating that S10 binding does not release the bound RNAP.

We conclude that the S10 interaction site of NusG-CTD is accessible in the NusG:RNAP complex and thus can promote ribosome binding and formation of a ribosome:NusG:RNAP complex.

To look for a ribosome:NusG-RNAP complex, we repeated the experiment using intact 70S ribosomes instead of S10^Δ^:NusB (Figure 3). In a first test, we titrated [ILV]-NusG with 70S ribosomes (Figure 3A). As in the [ILV]-NusG:RNAP experiment, signal intensity of [ILV]-NusG methyl groups was significantly, but not uniformly, decreased. In the presence of a twofold molar excess of ribosomes some NusG-NTD signals remained visible, whereas most NusG-CTD signals were nearly completely extinguished. Quantitative analysis of the [ILV]-NusG methyl group signal intensity in the presence of 0.5 equivalents of 70S ribosomes clearly shows that the relative intensity of NusG-CTD signals was in a narrow range < 2 %, whereas the relative intensity of NusG-NTD signals covered values from 0-4 %, and was higher on average (Figure 3B). Relative intensities of zero of NusG-NTD signals can be attributed to the fact that these signals are weak even in free NusG, and can thus not be quantified upon ribosome binding. Due to the flexibility of the linker, signals corresponding to amino acids in this region had the highest relative signal intensities. From these results we conclude that NusG binds to the ribosome via its CTD, in agreement with our cryo-EM structure (Figure 1). Due to the drastic increase in molecular mass we were unable to determine a binding site from these experiments, but nevertheless, the pattern of intensity changes of NusG-CTD signals was similar to that resulting form the titration of RNAP-bound NusG with S10, i.e. the most drastic decrease of signal intensity can be observed for residues 160-170, which are part of β-strands 3 and 4 and the intervening loop. Consequently, we conclude that the ribosome binding site is identical with the binding site for isolated S10.

**Figure 3:**
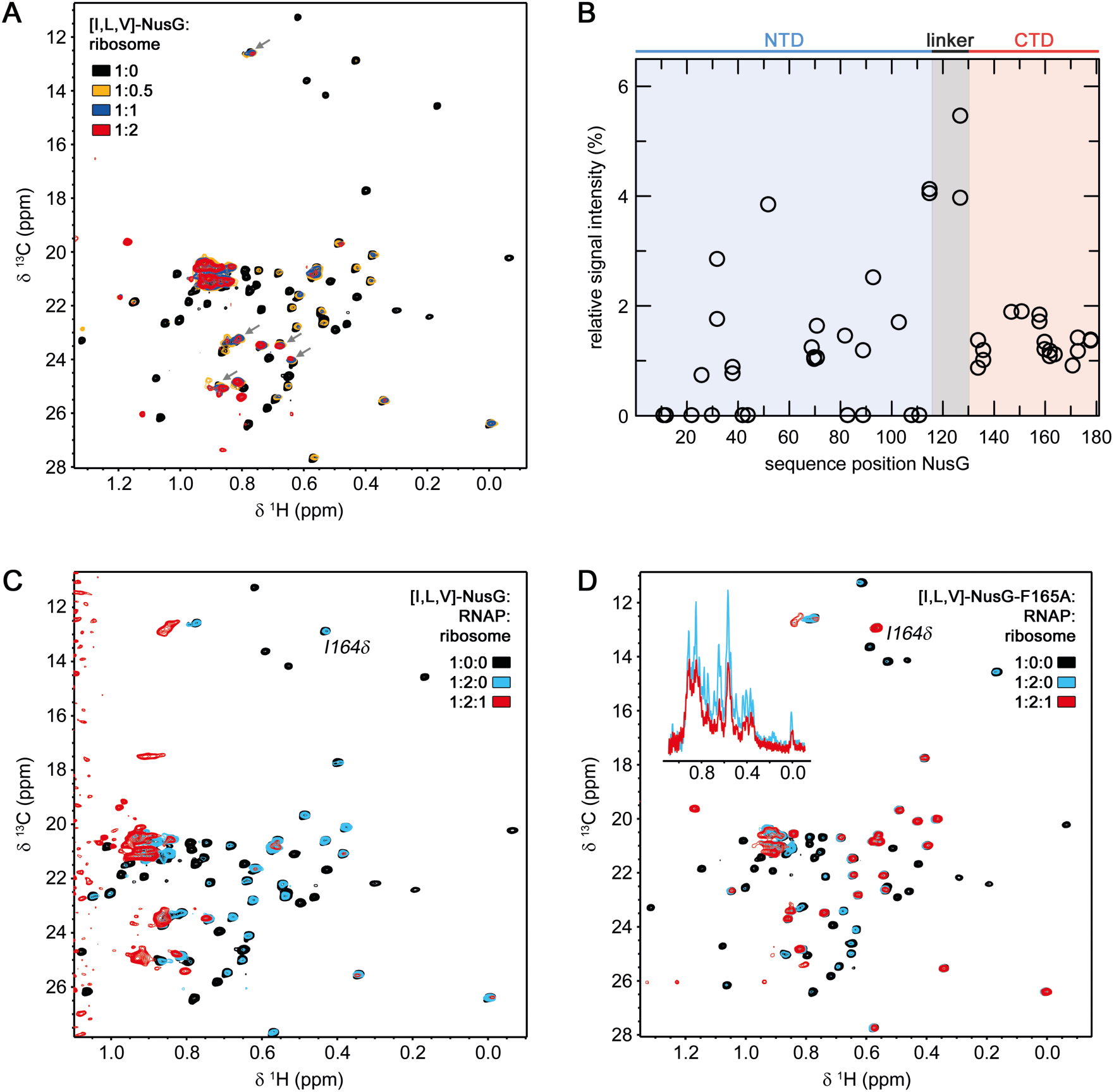
RNAP-bound NusG interacts with the 70 S ribosome. (A,B) NusG interacts with 70S ribosome via its CTD. **(A)** 2D [^1^H, ^13^C]-methyl-TROSY spectra of free [ILV]-NusG (11 µM, black) and [ILV]-NusG in the presence of 70S ribosome (molar ratio [ILV]-NusG:ribosome = 1:0.5 (6.6 µM [ILV]-NusG, orange); = 1:1 (7.5 µM [ILV]-NusG, blue); =1:2 (4 µM [ILV]-NusG, red). Arrows indicate [ILV]-NusG-NTD signals that are well visible in the [ILV]-NusG:ribosome complex. **(B)** Quantitative analysis of [ILV]-NusG methyl group signal intensities in the presence of 0.5 equivalents of 70S ribosome. Relative signal intensities are plotted versus the sequence position of NusG. The domain organization of NusG is indicated above the diagram. **(C)** 2D [^1^H, ^13^C]-methyl-TROSY spectra of [ILV]-NusG (11 µM, black), [ILV]-NusG in the presence of RNAP (molar ratio 1:2, 6 µM [ILV]-NusG, blue), and [ILV]-NusG in the presence of RNAP and 70S ribosome (molar ratio 1:2:1, 6 µM [ILV]-NusG, red). **(D)** 2D [^1^H, ^13^C]-methyl-TROSY spectra of [ILV]-NusG^F165A^ (20 µM, black), [ILV]-NusG^F165A^ in the presence of RNAP (molar ratio 1:2, 6 µM [ILV]-NusG^F165A^, blue), and [ILV]-NusG^F165A^ in the presence of RNAP and 70S ribosome (molar ratio 1:2:1, 6 µM [ILV]-NusG^F165A^, red). The inset shows the normalized 1D spectra of the corresponding titration step.

Next, we formed a complex of [ILV]-NusG and RNAP (molar ratio 1:2). The 2D methyl-TROSY spectrum of the complex revealed a decrease of signal intensities typical for NusG binding to RNAP (see Fig. 2C), i.e. primarily NusG-CTD signals remained visible. When we then added one equivalent of 70S ribosomes nearly all [ILV]-NusG signals were diminished (e.g. the signal corresponding to I164, which is in the loop responsible for ribosome binding). Strikingly, the spectrum differs from the spectrum of [ILV]-NusG in the presence of 70S ribosome (Fig. 3A). These results can be explained by three scenarios: (i) NusG-NTD is bound to RNAP, NusG-CTD is bound to a ribosome, and the ribosome directly interacts with RNAP, (ii) NusG-NTD is bound to RNAP, NusG-CTD is bound to the ribosome, but the ribosome does not interact with RNAP, (iii) NusG-NTD is bound to RNAP, the ribosome directly interacts with RNAP, and NusG-CTD is free, but is in the vicinity of the ribosome. To exclude the last scenario we repeated the experiment using a NusG variant, NusG^F165A^, in which F165, essential for ribosome binding (Burmann et al., 2010; Knowlton et al., 2003), is substituted by an Ala. Having ensured that the amino acid substitution does not influence the structure of NusG (Fig. 3-figure supplement 1A) we tested in a control experiment [ILV]-NusG^F165A^ binding to S10^Δ^. Indeed, we could detect no interaction (Fig. 3-figure supplement 1B,C). When we added 70S ribosomes to a preformed [ILV]-NusG^F165A^:RNAP complex (molar ratio 2:1), the spectrum corresponding to the [ILV]-NusG^F165A^:RNAP complex did not significantly change and, in particular, NusG-CTD signals remained visible, suggesting that the ribosome was not bound. However the general decrease in signal intensity indicated a direct RNAP:ribosome interaction. Thus, we conclude that NusG can serve as physical linker between ribosome and RNAP, although it remains elusive if a direct interaction between RNAP and a ribosome occurs in this NusG-coupled complex.

### Translation promotes NusG attachment to TEC

Chromatin Immuno Precipitation (ChIP) analysis showed that NusG binds to TEC well after transcription and translation initiation (Mooney et al., 2009b). Thus, we asked whether translation was, in fact, required for attachment of NusG to TEC. To approach this question, we examined the effects of translation on NusG-mediated Rho-dependent termination within the *lac* operon (Fig. 4A, Table 1) as NusG recruitment to the TEC is necessary for efficient Rho-dependent termination. Rho-dependent termination occurs within *lacZ* both *in vitro* (Burns and Richardson, 1995) and, upon the introduction of *lacZ* nonsense mutations, *in vivo* (Adhya and Gottesman, 1978; Newton et al., 1965). Polarity was measured using a probe to *lacA*, comparing mRNA levels with or without treatment with the Rho inhibitor bicyclomycin (BCM). Wildtype (WT) cells revealed no detectable termination (Table 1, Fig. 4A-I), which may be attributed to (i) sequestering of NusG-CTD by the ribosome, (ii) binding of the ribosome to the nascent RNA, or (iii) both. In all scenarios, however, the presence of the translating ribosome prevents Rho binding. We interfered with translation initiation by mutating the ribosome-binding site, i.e. the Shine-Dalgarno (SD) sequence (Fig. 4A-II), or translation elongation by introducing six successive rare arginine codons at two different locations in *lacZ* (Fig. 4A III, IV). Introduction of two G to A mutations in the *lacZ* SD sequence prohibits translation initiation of *lacZ* (Fig. 4A-II). *lacA* mRNA measurements gave a read-through of 21%, indicating that Rho-dependent termination occurs, but was inefficient in the absence of translation of *lacZ* mRNA. Introduction of the six in-frame rare arginine residues at the +4 position of *lacZ* (Fig. 4A-III, Table 1) allowed 29% read-through, i.e. Rho-dependent termination is present, but still inefficient if translation of *lacZ* mRNA is interfered with at early elongation. In contrast, introduction of the rare arginine residues 200 nt from the start site of transcription (Fig. 4A-IV, Table 1) resulted in high polarity, yielding < 1 % read-through. As efficient Rho-dependent termination requires NusG our results suggests that NusG binding to TEC occurs late and is dependent on translation.

**Figure 4:**
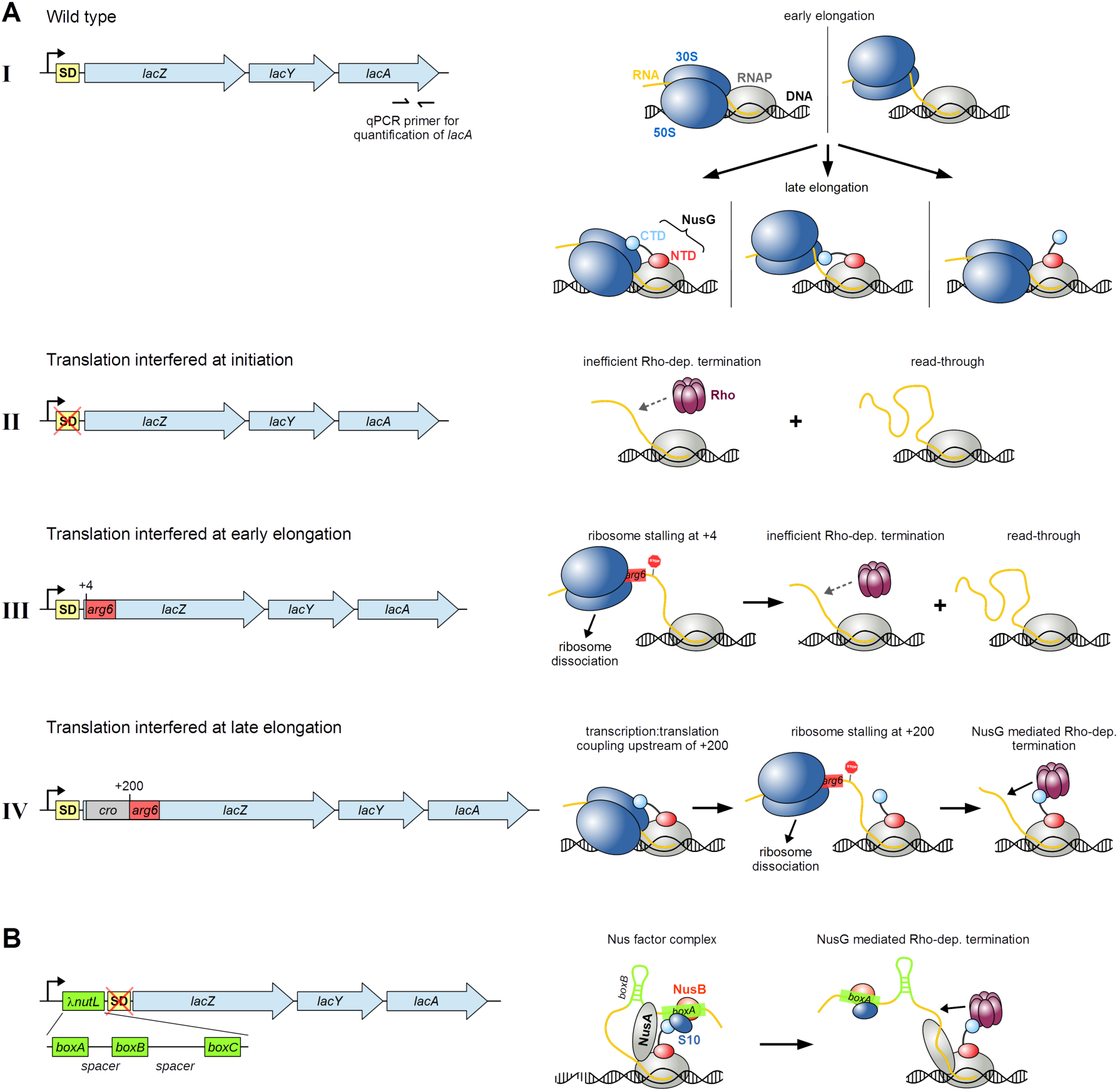
Translation is required for NusG recruitment to the TEC. (A, B) Left: Organization of the *E. coli lac* operon in strains MDS42 (A-I; wild type *lacZ*), RSW1225 (A-II; mutant (inactive) *lacZ* SD sequence), RSW1245 (A-III; in-frame insertion of six rare Arg codons (*arg6*) at position +4 of *lacZ*), RSW1276 (A-IV; in-frame insertion of λ*cro* and six rare Arg codons at position +4 of *lacZ* (equivalent to *arg6* being at position +200 of the gene)), and RSW1297 (B; λ*nutL* site upstream of mutant *lacZ* SD sequence). SD sequences of *lacY* and *lacA* were omitted for clarity. qPCR primers specific to the 3’ end of *lacA* (position indicated in A-I) were used to measure mRNA levels and thereby read-through of *lacA* (see Table 1). Right: Schemes of possible effects on transcription:translation coupling and Rho-dependent termination within *lacZ*. A-I, top: Ribosomes are recruited in the early elongation phase, leading to a directly coupled RNAP:ribosome complex (left) or uncoupled transcription and translation (right). A-I, bottom: NusG is recruited in late elongation, resulting in a NusG-coupled complex with (left) or without direct RNAP:ribosome contacts (middle), or modifying the pre-existing RNAP:ribosome complex without establishing an CTD:S10 interaction (right). A-II: Failure of NusG recruitment results in inefficient Rho-dependent termination and high *lacA* read-through. A-III: *arg6* stops the translating ribosome at position +4, whereas transcription elongation proceeds (left), resulting in ribosome dissociation and no NusG recruitment. Transcription proceeds and is only inefficiently terminated by Rho (right). A-IV: NusG couples transcription and translation (left) until *arg6* stops the ribosome at position +200 (middle), allowing efficient, NusG-stimulated Rho-dependent termination (right). B: λ*nutL* recruits NusA, NusG and the S10/NusB dimer, creating a Nus complex (left). NusG can thus support Rho-dependent termination.

**Table 1.** NusG couples late after transcription initiation. β-galactosidase was induced for 20 min from the *lac* operon with1mM IPTG. Where indicated, Rho-dependent termination was inhibited by adding 100μg/ml BCM 1 minute prior to induction. Read-through was calculated from the fold-increase of *lacA RNA* compared to *ompA* RNA in the presence or absence of BCM. RNA levels were measured using qRT-PCR and the fold-increase calculated using the ΔΔC_t_ method (Livak and Schmittgen, 2001). RSW1225 carries two G to A mutations in the *lacZ* ribosome-binding site. RSW1245 carries an insertion of six rare arginine codons (atg-acc-atg-AGG-AGA-CGA-AGG-AGA-CGA) at the amino terminus of *lacZ*. RSW1276 contains six rare arginine codons 200nt distal to the start of translation. RSW1297 carries an insertion of λ*nutL* immediately 5’ to the mutated ribosome binding site.

| strain | *lacZ* | *nutL* | BCM<sup>-</sup> | BCM<sup>+</sup> | RT (%) |
| --- | --- | --- | --- | --- | --- |
| MDS42 | wt | - | .25 | .26 | 99 |
| RSW1225 | SD<sup>-</sup> | - | .12 | .56 | 21 |
| RSW1245 | arg(6) - early | - | .13 | .49 | 29 |
| RSW1276 | arg(6) - late | - | <.01 | .12 | <1 |
| RSW1297 | SD<sup>-</sup> | + | .01 | .59 | 1 |

To confirm the hypothesis that NusG failed to attach to TEC in the absence of translation, we asked if a complex comprising Nus factors A, B, and E (Nus complex) assembled at a λ *nutL* site was able to recruit NusG so that it associates with TEC. Accordingly, we introduced the λ *nutL* site just upstream of the flawed *lacZ* SD sequence and measured *lacA* mRNA (Figure 4B, Table 1). Indeed, Rho-dependent termination was highly efficient, indicating that NusG had been recruited to TEC. Thus, counterintuitively, the Nus complex, which normally suppresses transcription termination in ribosomal (*rrn*) operons and, together with λN, on the phage λ chromosome, stimulates termination in this case.

We finally demonstrated that reduced termination efficiency in the mutant with the non-functional SD sequence was due to the failure of NusG recruitment to the TEC. In this assay we monitored Rho-dependent termination in a fusion construct that carries λ *cro,* the λ *nutR* site, the Rho-dependent λ *tR1* terminator and a *lacZ* reporter, with *lacZ* expression being heat-inducible (Fig. 5). Termination at the λ *tR1* site is poor when *cro* is translated, as seen with the *cro ms27* fusion (Table 2A, Fig. 5A-I; in the presence of an intact SD sequence we used *cro ms27*, where codon 27 carries a missense mutation so that the resulting protein is non-functional). The 3’ end of *cro* is adjacent to the λ *tR1* terminator, limiting the amount of free RNA available for Rho attachment if *cro* mRNA is translated. When λ *cro* carried a SD mutation translation initiation was ablated, but nevertheless there was significant termination at λ *tR1* (Table 2A, Figure 5A-II, compare read-through values with and without BCM). Formation of the Nus complex at λ *nutR* allows NusG recruitment and efficient termination. In the absence of NusB, the complex does not assemble, and there is extensive read-through at λ *tR1*.

**Figure 5:**
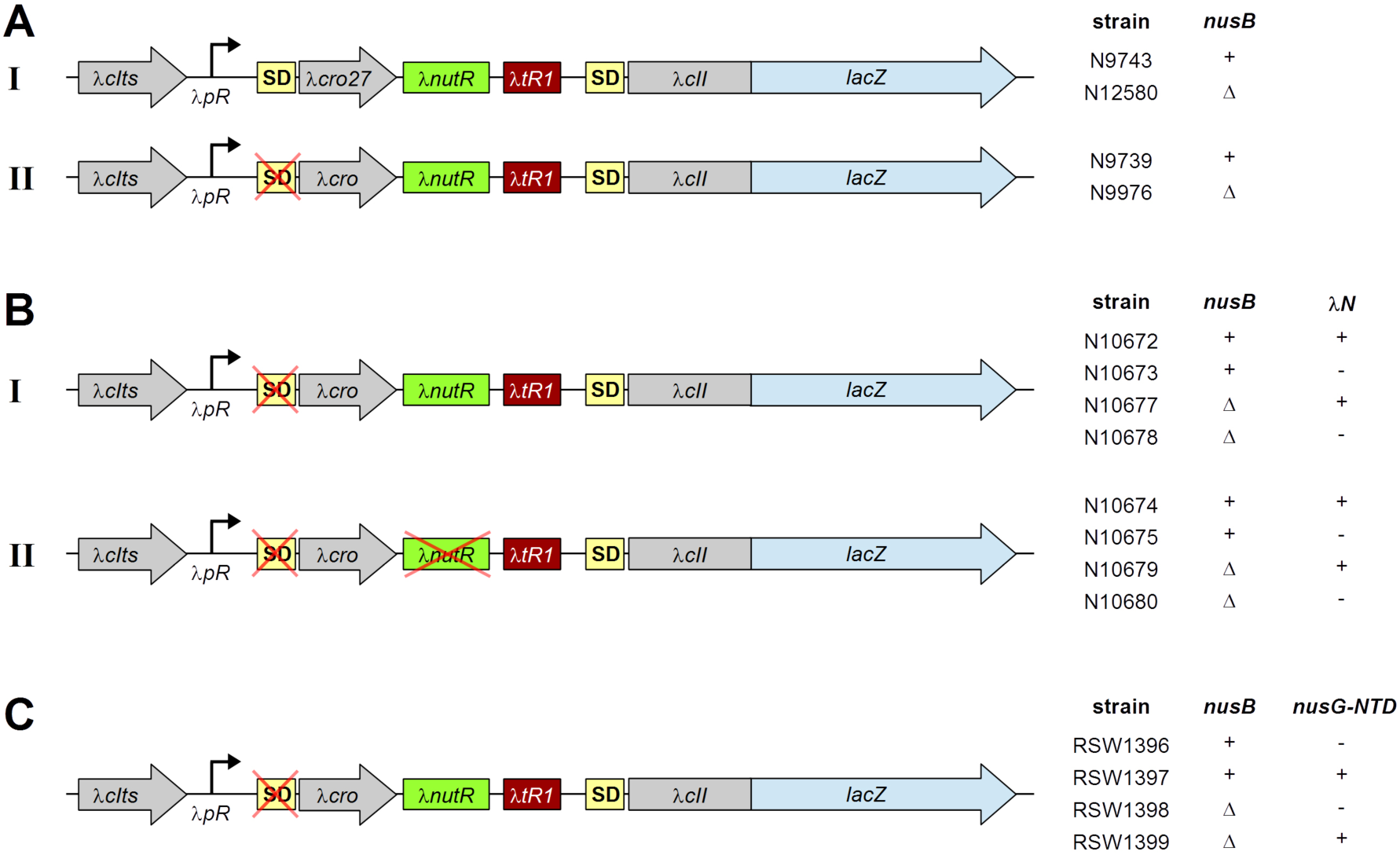
NusG can be recruited *via* a Nus complex. Genetic constructs used to monitor NusG mediated Rho-dependent termination are shown with the corresponding strains and their properties indicated on the right side. Transcription is started from the λ*pR* promoter, followed by WT-λ*cro* or λ*cro* carrying a missense mutation at codon 27 (λ*cro27*), a WT or mutant λ*nutR* site (B), the Rho-dependent terminator λ*tR1* and a λ*cII::lacZ* transcriptional fusion with corresponding SD site. All strains encode a temperature sensitive λcI construct (λ*cIts*) to allow temperature-controlled induction of gene expression from the λ*pR* promoter. λ*N^+^* strains listed in (B) further encode the λN protein; NusG-NTD for strains listed in (C) was supplied from plasmid pRM442. See also tables 2A-C.

**Table 2.** NusG recruitment depends on translation. (A. ) NusG coupling at *nutR* requires NusB. Expression of β-galactosidase was induced from a chromosomal *cII*::lacZ transcriptional fusion (λ*cIts-pR-cro-nutR-tR1-cII::lacZ*) by incubating at 42^0^ C for 30 min. N9743 and N12580 carry a missense mutation at *cro* codon 27, N9739 and 9976 have a G to C mutation in the *cro* Shine Dalgarno sequence (*SD*-), N12580 and N9976 are deleted for *nusB*. Where indicated, bicyclomycin (BCM) was added to 100μg/ml prior to induction of β-galactosidase. Read-through (RT) was calculated from the ratio of β-galactosidase activity in the presence or absence of BCM. **(B)** BoxA mutations block NusG coupling at *nutR*. Expression of β-galactosidase was induced from a chromosomal *cII*::lacZ transcriptional fusion (λ*cIts-pR-cro* (*SD^-^*) *-nutR-tR1-cII::lacZ*) by incubating at 42^0^ C for 30 min. Strains N10672, N10674, N10677 and N10679 express the λN transcription termination inhibitor. *boxA69* and Δ*nusB* strain numbers are indicated in Table 3. Read-through (RT) was calculated from the ratio of β-galactosidase activity in the presence or absence of λN. **(C)** NusG-NTD uncouples. Expression of β-galactosidase was induced from a chromosomal *cII*::lacZ transcriptional fusion (λ*cIts-pR-cro(*SD^-^*)-nutR-tR1-cII::lacZ*) by incubating at 42^0^ C for 30 min. The NusG-NTD was induced from the plasmid pRM442 *tac* promoter with 1mM IPTG for 10 min prior to induction of β−galactosidase in strains RSW1397 and RSW 1399. Strains RSW1396 and RSW1398 carried an empty vector (p*trc99A*) and were exposed to IPTG as above. Where indicated bicyclomycin (BCM) was added to 100μg/ml prior to induction of β-galactosidase. Read-through (RT) was calculated from the ratio of β-galactosidase activity in the presence or absence of BCM.

| **A** | **strain** | ***cro*** | ***nusB*** | **BCM<sup>-</sup>** | **BCM<sup>+</sup>** | **RT (%)** |
| --- | --- | --- | --- | --- | --- | --- |
|  | 9743 | *ms27* | + | 530 | 680 | 78 |
|  | 12580 | *ms27* | D | 890 | 1150 | 78 |
|  | 9739 | SD- | + | 141 | 613 | **23** |
|  | 9976 | SD- | D | 1191 | 1290 | 92 |

| **B** | **strain** | ***boxA*** | ***nusB*** | **λN<sup>-</sup>** | **λN<sup>+</sup>** | **RT (%)** |
| --- | --- | --- | --- | --- | --- | --- |
|  | 10673,<br>10672 | + | + | 126 | 946 | **13** |
|  | 10675,<br>10674 | *69* | + | 1212 | 2211 | 55 |
|  | 10678,<br>10677 | + | D | 2874 | 2616 | 100 |
|  | 10680,<br>10679 | *69* | D | 1896 | 2416 | 78 |

| **C** | **strain** | ***nusG*-**<br>***NTD*** | ***nusB*** | **BCM<sup>-</sup>** | **BCM<sup>+</sup>** | **RT (%)** |
| --- | --- | --- | --- | --- | --- | --- |
|  | RSW1396 | - | + | 247 | 862 | **29** |
|  | RSW1397 | + | + | 944 | 1013 | 93 |
| | RSW1398 | – | $\Delta$ | 2013 | 2314 | 93 |
| | RSW1399 | + | $\Delta$ | 2360 | 2760 | 86 |

The *boxA69* mutation also reduces Nus complex formation at λ *nutR*, and like the *nusB*-mutation, enhances read-through of λ *tR1* (Table 2B, Fig. 5B). In this experiment, we suppressed termination at λ *tR1* with λ N antitermination factor instead of BCM. Finally, we showed that expression of *nusG-NTD*, which competes with NusG for binding to RNAP, enhances read-through (Table 2C, Fig. 5C). Taken together, these results strongly support the idea that NusG can be supplied by the Nus complex assembled at λ *nutR* in the absence of translation, inducing Rho-dependent termination at λ *tR1*.

### Discussion

We determined a cryo-EM structure of a NusG: 70S complex showing binding of one molecule NusG per ribosome, consistent with previous results (Saxena et al., 2018). NusG binds to the S10 protein on the 30S subunit via its CTD as indicated by the study of isolated NusG-CTD and S10 (Burmann et al., 2010); density for NusG-NTD was not observable, suggesting that it remains flexible. We must attribute the low occupancy of the NusG-CTD on the 70S ribosome in the cryo-EM experiment to weak binding adversely affected by the conditions of sample preparation. Notably, although tmRNA contacts the ribosome at various sites, the binding of NusG-CTD and tmRNA on S10 is mutually exclusive. This suggests a model in which uncoupling at rare codons, at which tmRNA releases ribosomes, is promoted by tmRNA-induced release of NusG (Roche and Sauer, 1999). The freed NusG:TEC complex exposes the NusG-CTD, and is then subject to Rho-dependent transcription termination.

Simultaneous binding of NusG to S10 and RNAP has been demonstrated by solution-state NMR studies, confirming the S10 binding site on NusG-CTD as identified in a binary NusG-CTD:S10 system (Fig. 2) (Burmann et al., 2010). Moreover, we show that NusG can bind RNAP and 70S ribosome simultaneously; this is the first direct structural evidence for NusG-mediated transcription:translation coupling. The flexibility of the linker between the NusG-NTD and the NusG-CTD permits these interactions.

The operon-specific *E. coli* NusG paralog, RfaH, likewise simultaneously binds S10 and RNAP in the context of a paused TEC (Burmann et al., 2012; Zuber et al., 2019). RfaH, which also comprises an NTD and a flexibly-connected CTD (Belogurov et al., 2007; Burmann et al., 2012), uses the same binding sites as NusG to interact with RNAP and S10 (Burmann et al., 2010, 2012; Kang et al., 2018; Sevostyanova et al., 2011; Zuber et al., 2019). However, RfaH, unlike NusG, complexes with TEC early after transcription initiation, when TEC pauses at an operon polarity suppressor (*ops*) site, a representative of the *E. coli* consensus pause sequence (Larson et al., 2014; Vvedenskaya et al., 2014). Located in the untranslated leader region of RfaH-controlled operons, *ops* is responsible for sequence-specific recruitment of RfaH (Zuber et al., 2018). Importantly, RfaH-dependent operons lack a consensus SD sequence. To initiate translation, RfaH recruits a ribosome to these mRNAs, making coupling essential for translation activation and efficient gene expression (Burmann et al., 2012). The binding modes of RfaH and NusG to RNAP and S10 are very similar, indicating that coupling as observed for RfaH can also be mediated by NusG and vice versa. However, once recruited, RfaH excludes NusG (Kang et al., 2018), thus preventing intra-operon Rho-dependent transcription termination in RfaH-controlled operons (see (Artsimovitch and Knauer, 2019)).

We have confirmed the results of Mooney et al that NusG binds to TEC only after significant RNA synthesis (Mooney et al., 2009b). As postulated by these authors, binding depends on active translation of the mRNA. Thus efficient Rho-dependent transcription termination, which requires the attachment of Rho to the NusG-CTD, does not occur at the end of an untranslated gene. We have shown that the failure of NusG to bind TEC is responsible for the absence of termination. Thus, placing a λ *nut* site at the start of the gene recruits NusG and restores termination. At present, it is not understood why NusG appears to be delivered to TEC by ribosomes *in vivo*, whereas it binds directly to RNAP in a purified system lacking ribosomes. A possible explanation would be that NusG binds to RNAP discontinuously in an on-and-off mode in the untranslated leader region and that the NusG:RNAP interaction is only stabilized when the ribosome is coupled upon translation initiation. We should recall that NusG has two binding sites in the coupled system, which significantly increases its affinity.

A direct connection between transcription and translation was first predicted in 1964 (Byrne et al., 1964). Transcription:translation coupling is necessary to coordinate gene expression and to maintain genome stability (McGary and Nudler, 2013). In 1970, Miller *et al*. performed electron miscroscopy analyses of lysed *E. coli* cells (Miller et al., 1970). They demonstrated that all mRNA molecules are connected to the *E. coli* genome, and that the ribosome at the newly synthesized end of a polyribosome is almost always immediately adjacent to the putative RNAP molecule. They concluded that translation is completely coupled with transcription. Coupling could allow RNAP to monitor the translation rate while providing newly synthesized mRNA to the ribosome. The structural basis of this coupling is, however, still only poorly understood. Our results strongly suggest that NusG may mediate coupling. However, since NusG attaches to the TEC downstream to the translation initiation site, the coupled transcription:translation complex must initially consist of a ribosome bound directly to TEC, in agreement with two cryo-EM structures and *in vitro* data (Demo et al., 2017; Fan et al., 2017; Kohler et al., 2017). One cryo-EM structure shows the so-called “expressome”, an RNAP:70S complex generated by letting a ribosome translate until it encounters a stalled RNAP (Fig. 6A; (Kohler et al., 2017)). In this complex RNAP directly binds to the 30S subunit with the RNA exit region of RNAP docking onto the ribosome near the mRNA tunnel entry between ribosomal proteins S3, S4, and S5. In this structure, mRNA exiting from RNAP can directly enter the ribosome. Another cryo-EM structure showed an RNAP:30S complex generated by mixing 30S subunit with a 3-fold excess of RNAP (Fig. 6B; (Demo et al., 2017)). In this structure RNAP is bound to the 30S subunit near the mRNA binding site between the head and the platform domains, contacting ribosomal proteins S1, S2, S18, S21, and hairpin loop 40 of 16S rRNA, in agreement with crosslinking data (Fan et al., 2017). Strikingly, this position is located more than 80 Å from the binding site observed in the expressome structure, i.e. on the opposite side of the 30S head. Importantly, it ensures that RNAP interacts with the cytosolic site of the 30S ribosomal subunit so that the nascent RNA exiting from RNAP is directly guided to the entry site on the ribosome. Assuming that the RNAP:30S complex corresponds to a coupling complex at translation initiation and the expressome structure to a translation elongation complex, a would require a massive relocalization of RNAP.

**Figure 6:**
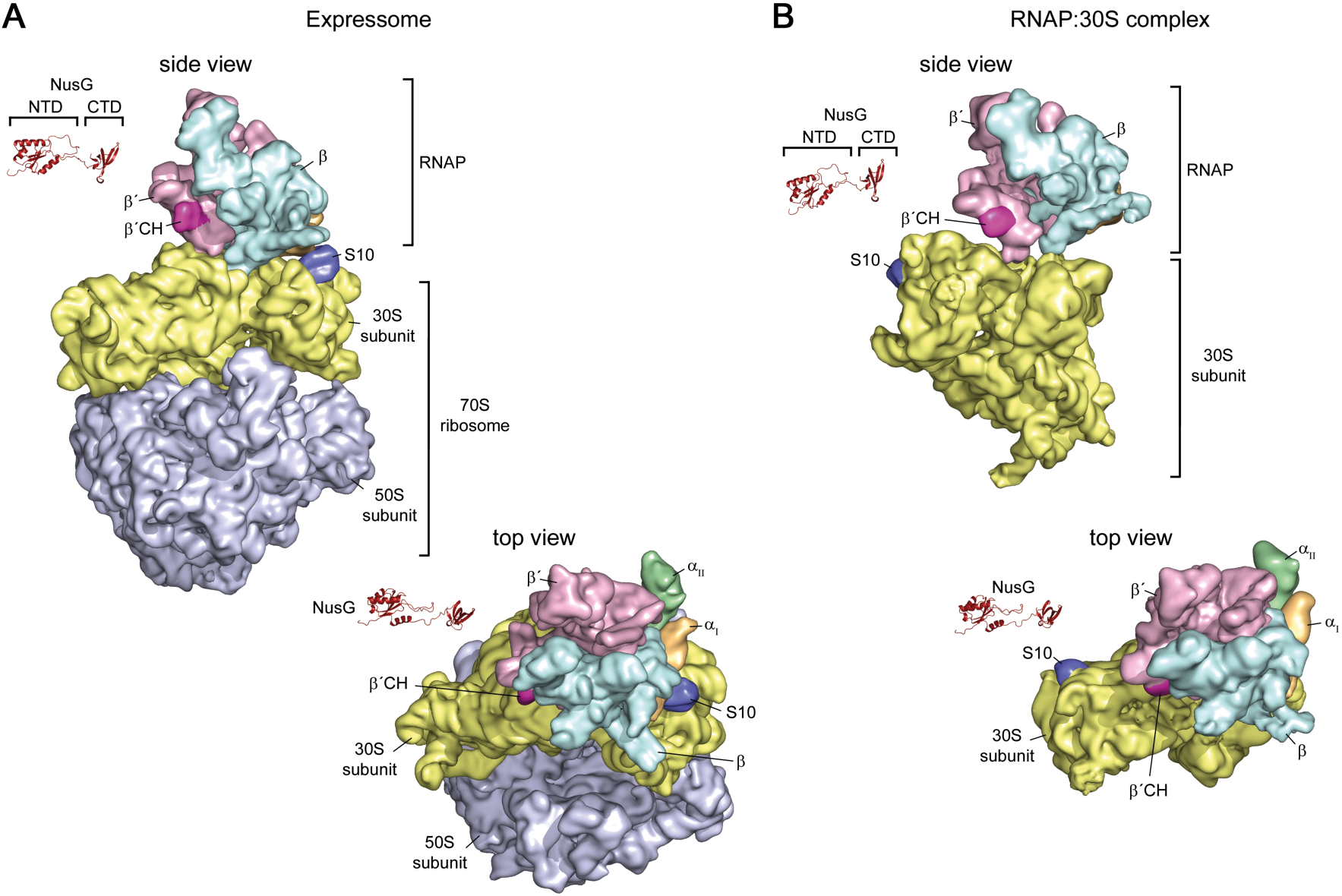
Structures of coupling complexes. Structures of the expressome (A) and an RNAP:30S complex (B) determined by cryo-EM are shown. RNAP and ribosomal subunits are in surface representation, NusG is shown as ribbon. α_I_, orange; α_II_ green; β, cyan; β’, light violet; 30S, yellow; 50S, light blue; β’CH, pink; S10, dark blue. PDB IDs: expressome, 5MY1 and 6O9J; RNAP:30S, 6AWD; NusG-NTD: 2K06; NusG-CTD: 2JVV

Interestingly, neither the RNAP:30S nor the expressome structures allow NusG-(or RfaH-) mediated coupling: the linker of NusG/RfaH is too short (Fig. 6). However, as the cryo-EM structures suggest that the position of RNAP on the 30S subunit might be flexible, these structures could be snapshots of distinct situation during translation. Thus, we suggest that at some distance downstream of the translation initiation site, NusG recognizes and enters the coupling complex.

In summary, we hypothesize that two coupling modes exist, a direct coupling between the ribosome and TEC during translation initiation and early elongation and a NusG-mediated coupling mode later in translation. The question of whether the 70S ribosome still directly contacts TEC in the NusG-mediated system remains elusive. The expressome structure (Kohler et al., 2017) does not allow simultaneous NusG binding to TEC and the 70S ribosome, thus the relative orientation of 70S ribosome to TEC might be different in the direct and the NusG-mediated system. The latter may thus require a reorientation of the TEC and the 70S ribosome and confer the system more flexibility, necessary to keep transcription and translation synchronized, even if these processes are differently regulated or occur at different rates.

## Methods

### Strain Construction

Standard bacteriological techniques used in strain construction (e.g., transformation, transduction and media preparation) are as described in Silhavy et al. (1984). Standard molecular biology techniques were as described in Sambrook and Russell (Sambrook and Russel, 2001). N10780 was constructed by P1 transduction of *rpoC-his:kanR nusGF165A* from NB885 into MDS42. N11158 was constructed by P1 transduction of Δ*ssrA::camR* from RSW943 into MDS42. N11816 was constructed by P1 transduction of Δ*relA::kanR* from RLG847 into N11158. RSW1008 was constructed by P1 transduction of Δ*ssrA::camR* from RSW943 into N4837. RSW1010 was constructed by P1 transduction of *rpoC-his:kanR nusGF165A* from NB885 into N4837. RSW1012 was constructed by P1 transduction of Δ*ssrA::camR* from RSW943 into RSW1010. RSW1175 was constructed by P1 transduction of *ΔrelA::kanR* and *ΔspoT::camR* from RLG847 into MDS42. RSW1245 was generated using recombineering (Sharan et al., 2009) to introduce six rare arginine codons (atg-acc-atg-AGG-AGA-CGA-AGG-AGA-CGA-att-acg-gat) into the 5’end of *lacZ* in MDS42 changing the amino acid sequence of the aminoterminus from MTMITD to MTMRRRRRRITD with six inefficiently translated arginine codons. RSW1225 was produced using recombineering to introduce two G to A mutations in the ribosome binding site of *lacZ* in MDS42. This resulted in a change from…TTCACACA<u>GG</u>AAACAGCTatgaccatg… to …TTCACACA<u>CC</u> AAACAGCTatgaccatg…inactivating the ribosome binding site. RSW1225 is *lac^-^*.

### Cloning

The plasmid encoding NusG-F165A was generated by site-directed mutagenesis according to the QuikChange Site-Directed Mutagenesis Kit (Stratagene), using vector pET11A_*nusG* (Burmann et al., 2011) as template and primers Fw_NusG-F165A (5’ GTG TCT GTT TCT ATC GCG GGT CGT GCG ACC CCG 3’) and Rv_NusG-F165A (5’ CGG GGT CGC ACG ACC CGC GAT AGA AAC AGA CAC 3’; both primers were obtained from metabion, Martinsried, Germany).

### Protein production and isotopic labeling

NusG and NusG-NTD were produced as described (Burmann et al., 2011) as were NusG-CTD (Burmann et al., 2010) and RNAP and S10^Δ^:NusB used for NMR spectroscopy (Zuber et al., 2019). Production of NusG-F165A was analogous to NusG (Burmann et al., 2011). For unlabeled proteins, bacteria were grown in lysogeny broth (LB) medium. [^1^H,^13^C]-labeling of methyl groups of Ile, Leu, and Val residues in perdeuterated proteins was accomplished by growing bacteria in M9 medium (Meyer and Schlegel, 1983; Sambrook and Russel, 2001) prepared with increasing amounts of D_2_O (0 % (v/v), 50 % (v/v), 100 % (v/v); Eurisotop, Saint-Aubin, France) and (^15^NH_4_)_2_SO_4_ (CortecNet, Voisins-Le-Bretonneux, France) and d_7_-glucose (Cambridge Isotope Laboratories, Inc., Tewksburgy, USA) as sole nitrogen and carbon sources, respectively. Amino acid precursors (60 mg/l 2-keto-3-d_3_-4-^13^C-butyrate and 100 mg/l 2-keto-3-methyl-d_3_-3-d_1_-4-^13^C-butyrate; Eurisotop, Saint-Aubin, France) were added 1 h prior to induction. Expression and purification protocols were identical to those of the non-labeled proteins.

Intact 70S ribosomes were produced as follows. *E. coli* strain MRE600 cells grown in LB medium were harvested, lysed by passing through the French Press 3x at ∼800 PSI, and clarified by a short centrifugation (20,000 rpm, 40min) in opening buffer (20 mM Tris-HCl pH=7.5, 100mM NH_4_Cl, 10.5 mM Mg acetate, 0.5 mM EDTA, with half a protease inhibitor cocktail tablet (Roche, EDTA-free), and 1mM TCEP added just before use). The lysate was loaded onto the top of 5 mL sucrose cushion (20 mM Tris-HCl, pH=7.5, 500 mM NH_4_Cl, 10.5 mM Mg acetate, 0.5 mM EDTA, 1.1 M sucrose, and 1 mM TCEP added before use) and centrifuged for 24 h at 28,000 rpm in a 70Ti rotor (Beckman Coulter, Inc.). The pellets were suspended in washing buffer (20 mM Tris-HCl, pH=7.5; 500mM NH_4_Cl, 10.5mM Mg acetate, 0.5mM EDTA and 1 mM TCEP added before use), and centrifuged through a 10–35% sucrose gradient for 19 h at 16,000 rpm in a SW28 rotor (Beckman Coulter, Inc.). Fractions containing the 70S peaks were pooled and kept at −80℃ for further use.

Ribosomes for NMR experiments were obtained from New England Biolabs.

### Electron Microscopy

Purified 70S ribosomes were incubated with full-length NusG at a ratio of 1:7 for 40 min at room temperature, prior to blotting and plunge-freezing as previously described (Grassucci et al., 2007). Data were collected on a TF30 Polara electron microscope (FEI, Portland, Oregon) at 300kV using a K2 Summit direct electron detector camera (Gatan, Pleasanton, CA). Images were recorded using the automated data collection system Leginon (Suloway et al., 2005) in counting mode, and taken at the nominal magnification of 32,000x, corresponding to a calibrated pixel size of 1.66Å.

### Image processing

A total of 188,127 particles were automatically extracted from 1327 images using Arachnid (Langlois et al., 2014). RELION (Scheres, 2012) 3D classification was used to resolve the heterogeneity of the particle images, and auto-refinement to further improve resolution for each class. The final refinement for the NusG-bound 70S class containing 17,122 particles yielded an average resolution of ∼6.8Å (FSC=0.143; following “gold standard” protocol).

### Model building

The starting model was assembled from the X-ray structure of the *E. coli* 30S ribosomal subunit (PDB ID 4GD2) and the NMR solution structure of the NusG-CTD (PDB 2KVQ chain G). This starting model was first docked into the segmented maps of our 70S density map as a rigid body using UCSF Chimera (Pettersen et al., 2004). Then it was fitted into the segmented map using the Molecular Dynamic Flexibly Fitting (MDFF) method (Trabuco et al., 2008) and run using the NAMD program (Phillips et al., 2005) for 0.5 ns of simulation time, followed by 5,000 steps of energy minimization.

### NMR spectroscopy

NMR experiments were conducted on Bruker Ascend Aeon 900 and 1000 MHz spectrometers equipped with cryogenically cooled, inverse triple resonance probes at 298 K. NMR data was converted and processed using in-house software. 2D correlation spectra were visualized and analyzed with NMRViewJ (One Moon Scientific, Inc., Westfield, NJ, USA), 1D spectra were plotted using MatLab (The MathWorks, Inc., Version 9.2.0.538062). Resonance assignments for NusG methyl groups were taken from a previous study (Mooney et al., 2009a).

[ILV]-NusG-CTD was in 10 mM K-phosphate (pH 7.5), 50 mM KCl, 1 mM MgCl_2_, 99.9% (v/v) D_2_O, [ILV]-NusG-NTD in 50 mM Na-phosphate (pH 7.5), 50 mM KCl, 0.3 mM EDTA, 5 % (v/v) d_7_-glycerol, 0.01 % (w/v) NaN_3_, 99.9 % D_2_O. For the titration of [ILV]-NusG with RNAP and S10^D^:NusB, all proteins were in 50 mM Na-phosphate (pH 7.5), 50 mM KCl, 0.3 mM EDTA, 99.9 % (v/v) D_2_O and 5 mM MgCl_2_ and 2 mM DTT were added to the NMR sample to increase the-long-term stability of RNAP. For all interaction studies involving ribosomes and for the titration of [ILV]-NusG-F165A with S10^Δ^:NusB, all components were in 20 mM HEPES-KOH (pH 7.6), 10 mM Mg-acetate, 30 mM KCl, 7 mM β-mercaptoethanol, 10 % D_2_O. The titration of [ILV]-NusG-F165A with S10^Δ^:NusB was conducted in a 5 mm tube with an initial sample volume of 550 µl. All other measurements were carried out in 3 mm NMR tubes with an (initial) volume of 200 µl.

1D and 2D titration experiments were evaluated quantitatively by analyzing either changes in signal intensity or changes in chemical shifts. If chemical shift changes were in the fast regime on the chemical shift the normalized chemical shift perturbation (Δδ_norm_) was calculated according to equation 1.

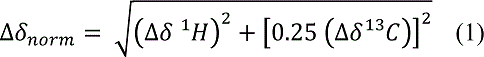

with Δδ being the resonance frequency difference between the initial and final state of the titration (i.e. [ILV]-NusG:RNAP:S10^D^:NusB = 1:2:0:0 *vs*. 1:2:2:2) in ppm.

If the system was in slow or intermediate chemical exchange the signal intensities were analyzed quantitatively. First, the intensity of each 1D spectrum or methyl group signal, respectively, was normalized by the concentration of the [ILV]-labeled protein, the receiver gain, the number of scans, and the length of the 90° ^1^H pulse. Then the relative intensity, i.e. the ratio of the normalized signal intensity of [ILV]-labeled protein in the respective titration step to the normalized signal intensity of free [ILV]-labeled protein, was calculated and plotted against the sequence of NusG or the NusG variant, respectively.

### qRT-PCR

Total RNA was extracted from cells grown in M9 medium supplemented with casamino acids (0.2%) at 37^0^C to mid-log phase (O.D. 600=0.3). Fold-increase of the PCR product was determined using qRT-PCR. cDNA was synthesized using RNA was extracted from logarithmically growing cultures (O.D._600_=0.2-0.3) Where indicated, cells were treated with BCM (100μg/ml) 1 min before induction with 1mM IPTG for *lacZ*. Samples were removed (0.5ml) at the indicated times and total RNA extracted RNA extracted using Qiagen RNeasy and Qiagen RNAprotect Bacteria Reagent (Qiagen, Germantown, MD). cDNA was synthesized from the samples using High Capacity RNA to cDNA kit (ThermoFisher, Waltham, MA). qRT-PCR reactions were performed using Taqman Gene Expression Master Mix (Thermofisher, Waltham, MA) and Biorad DNA Engine Opticon2 Real-Time Cycler (Bio-Rad Laboratories, Hercules, CA) and PrimeTime qPCR probes (Integrated DNA Technologies, Coralville, IA). The *lacA* transcript was probed with the following: probe:5’-/56-FAM/CCACATGAC/ZEN/TTCCGATCCAGACGTT/3IABkFQ/-3’; primer1:5’-ATACTACCCGCGCCAATAAC; primer2:5’-CCCTGTACACCATGAATTGAGA).

The reference gene was *ompA*(probe:5’-/56-FAM/CAACAACAT/ZEN/CGGTGACGCACACAC /3IABkFQ/-3’; primer1:5’-TGACCGAAACGGTAGGAAAC; primer2:5’-ACGCGATCACTCCTGAAATC). The PCR was performed using the following conditions: 50 ^°^C for 10 min., 95 ^°^C for 2 min, followed by 40 cycles each of 95 °C for 15s, and 60 °C 1min. All reactions were performed in duplicate. Fold increases were calculated from measured C_t_ values using the ΔΔC_t_ method (Livak and Schmittgen, 2001). Values are the average of three or more independent experiments.

### *β−*galactosidase assays

Cultures were grown in LB to early log phase (O.D. 600 =0.3) at 37 ^0^C. Where indicated bicyclomycin (BCM) (100μg/ml) was added to inhibit Rho-dependent transcription termination prior to induction of *lacZ* with 1mm IPTG. Where indicated λN was expressed by incubation at 42 ^°^C. Reactions were terminated 15 min. after induction. β-galactosidase was measured using a modification of the method of Miller (Zhang and Bremer, 1995). Readthrough was calculated from the ratio of β-galactosidase activity +/-BCM.

## Author Contributions

R.W., Y.H., M.S., W.L., B.S. P.K.Z. and S.H. performed the experiments. J.F., Y.H., M.S., R.W. P.K.Z. S.H.K. and M.G. designed the experiments and interpreted the results. M.S. M.G. M.S. P.K.Z. S.H.K. and J.F. wrote the paper.

## Acknowledgments

We gratefully acknowledge the help of D. Shapoval, M. Bubunenko, N. Costantino and D. Court. We have had numerous useful discussions with P. Roesch and his group, for which we are most indebted. Supported by HHMI and NIH R01 GM29169 (to J.F.), NIH R01 GM037219 (to M.G.) and the German Research Foundation grant Ro617/21-1 (to P.R.). S.H. was supported by an Amgen Fellowship.

---

**Figure 2-figure supplement 1:**
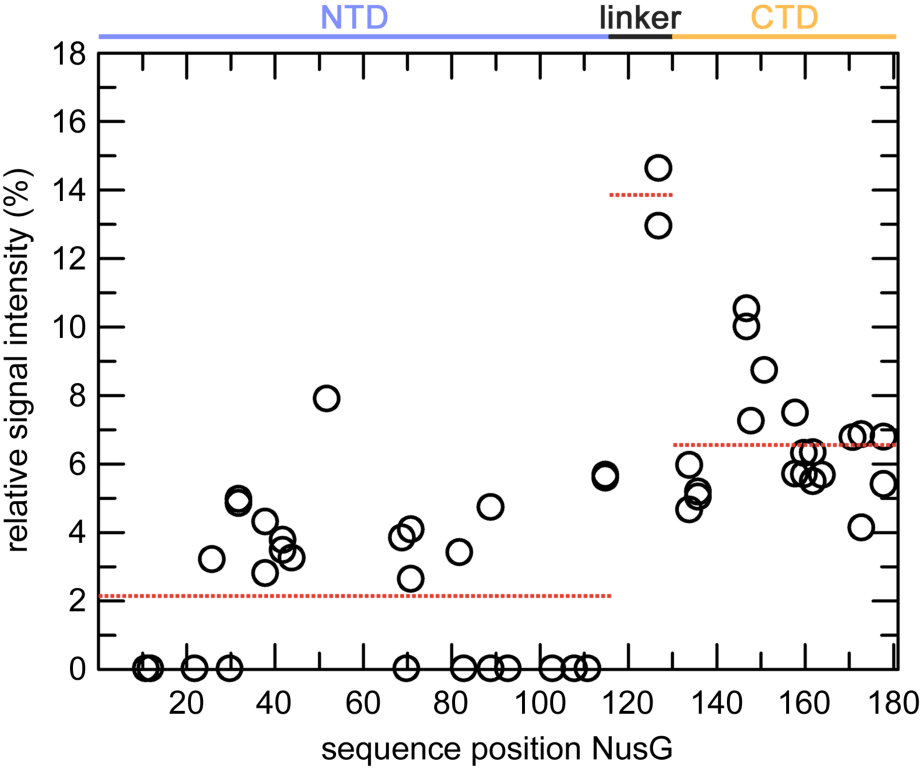
Binding of [ILV]-NusG to RNAP. [^1^H,^13^C]-methyl-TROSY derived relative signal intensities of [ILV]-NusG methyl groups after addition of two equivalents of RNAP (see Fig. 2B). Red, dashed horizontal lines indicate average relative signal intensities of NusG-NTD, the linker, and NusG-CTD (domain organization is indicated at the top). Related to Figure 2B.

**Figure 3-figure supplement 1:**
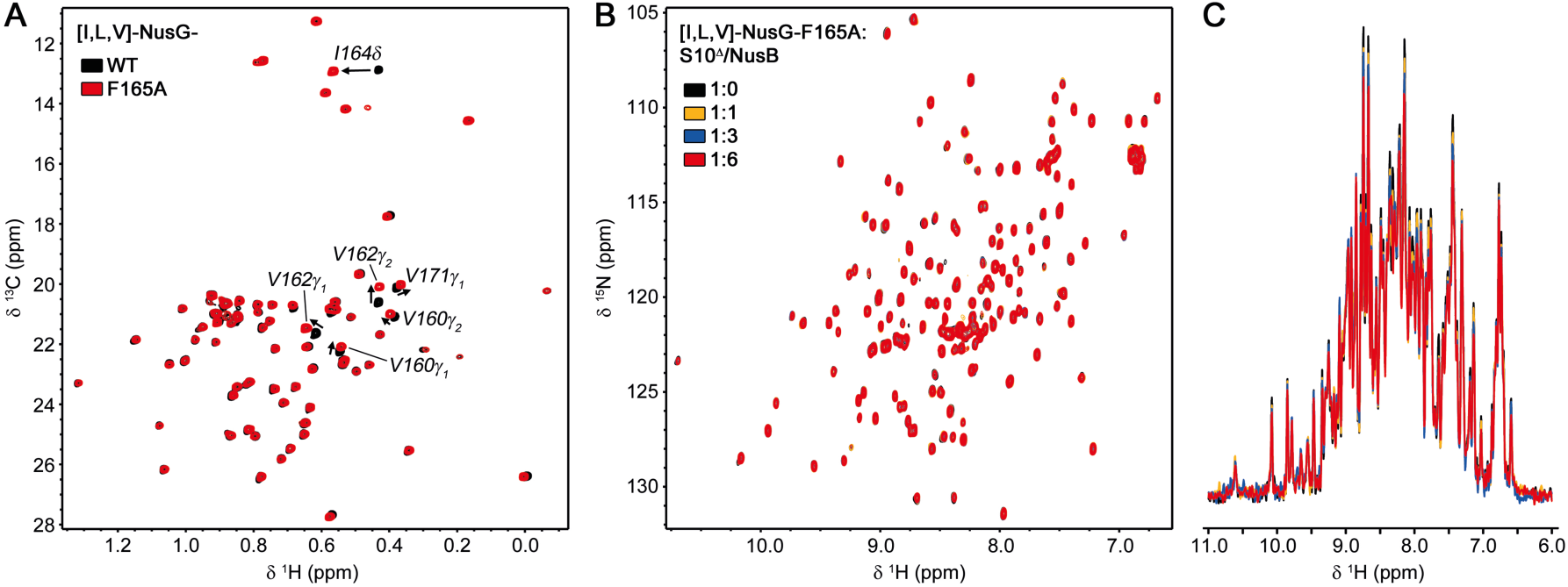
NusG-F165A does not interact with S10^Δ^/NusB. (A) 2D [^1^H, ^13^C]-methyl-TROSY spectra of [ILV]-NusG (11 µM, black) and [ILV]-NusG^F165A^ (20 µM, red). Arrows and labels indicate NusG-CTD methyl groups affected in their resonance frequencies by the F165A amino acid substitution. **(B,C)** 2D **(B)** and normalized 1D **(C)** [^1^H, ^15^N]-HSQC spectra of 20 µM [ILV]-NusG^F165A^ upon titration with 432 µM S10^Δ^/NusB (colors as indicated).

